# Free energy profiles of temozolomide crossing brain plasma membranes

**DOI:** 10.1101/2024.07.16.603685

**Authors:** Yanhong Ge, Huixia Lu, Jordi Marti

## Abstract

Temozolomide is an efficient small-molecule drug mostly employed for the treatment of glioblastoma, a tumor attacking both the spinal cord and the brain. Understanding the interactions of temozolomide with different lipids at the brain cell membrane can help elucidate how temozolomide permeates through cell membranes and its membrane-crossing ability. In the present work, we have constructed a simplified brain plasma membrane model to explore temozolomide’s microscopic structure and dynamics by means of all-atom microsecond scale molecular dynamics simulations. The preferential location of temozolomide is at the solvent-aqueous fluid surrounding the brain membrane, but it can access the interface with the membrane regularly, eventually binding to lipids of the choline and cerebroside classes. The free energy barriers of temozolomide related to brain-like plasma membrane crossing were investigated by adaptive biasing force methods, revealing values ranging from 18.5 to 66.5 kcal/mol at temperatures of 323 K and 310 K, respectively. Our results suggest that temozolomide cannot cross the membrane by pure diffusion at the normal human body temperature but that rising the temperature significantly increases the probability of barrier crossing. This fact is mainly due to the crucial role played by cholesterol and lipids of the cerebroside class. The findings reported in this work can be used to optimize the molecular design of temozolomide and to develop new analogs with better pharmacokinetic properties.

**Author summary:** Glioblastoma is a devastating tumor affecting the brain and spinal cord, which has in the FDA-approved drug temozolomide its main clinical treatment. The present study explores how temozolomide interacts with several lipids in brain-like cell membranes. Our findings show that at normal body temperature temozolomide cannot cross the membrane by pure diffusion, but that higher temperatures significantly enhance its ability to cross the membrane by reducing the free energy barriers. Temozolomide interacts differently with several lipids and sterols depending on the temperature, which affects its permeability. This implies that temozolomide will cross the outer layer of the brain membrane only with the help of driving agents, such as intermembrane proteins. Our research suggests that temozolomide may be more effective at higher temperatures and cancer patients with fever might need a lower dose. Importantly, cholesterol plays a key role in blocking temozolomide from crossing brain-like membranes, so reducing dietary intake of cholesterol and cerebroside lipids could help modify brain cell membranes, making it easier for temozolomide to target cancer cells effectively and potentially reducing side effects.

## Introduction

Temozolomide (TMZ) is the best known and efficient drug usually employed for patients suffering from glioblastoma, which is a malignant brain tumor of hard treatment [1, 2], and also for anaplastic astrocytoma [3]. TMZ is an imidazotetrazine small-molecule acting as an alkylating agent with remarkable efficiency for glioblastoma [4].

Nevertheless, some obstacles like the reticuloendothelial system and mainly the blood-brain barrier (BBB) [5] prevent most drugs from entering the brain and tumors inside. These obstacles are crucial to surmount in order to achieve the success in the treatment [6–8]. In normal conditions, TMZ with a short half-life in blood plasma (∼ 1.8 h) is totally absorbed in the gastrointestinal tract after oral administration [9].

Later on, the prodrug TMZ tends to hydrolyze rapidly into an active form of the unstable metabolite 5-(3-methyltriazen-1-yl) imidazole-4-carboxamide (MTIC, half-life ∼ 2 min) at physiologic pH [9–11], which rapidly degrades to 5-amino-imidazole-4-carboxamide and to the methyldiazonium ion [12]. As the final step, the methyldiazonium cation preferentially methylates DNA at guanine and adenine residues [13, 14]. It should be noticed that there is a quite narrow pH window near the physiological pH at which the whole process of TMZ activation can occur [15].

The MTIC compound produced in the plasma is not able to cross the BBB and is formed locally in the brain. However, TMZ is generally believed to penetrate the BBB relatively well [16]. As a result of prolonged therapy, severe and unpredictable myelosuppression effects precluded TMZ’s further development [17]. To increase the drug concentration of TMZ in the brain, many scientists believe that nanoscale delivery systems can be a promising strategy [11]. For instance, Bouzinab et al. [18] reported a nanodelivery system of biocompatible protein nanocage able to improve the delivery of TMZ to cancer cells, showing an enhanced potency of imidazotetrazine for the treatment of glioblastoma multiforme and wider-spectrum malignancies. Meanwhile, other scientific teams are concentrating on creating potential drugs, including derivatives of TMZ [19–21]. Recently Yin et al. [22] reported a new drug derived from TMZ and called 5-aminoimidazole-4-carboxamide (AICA).

The efficacy of TMZ is also reduced, due to its limited access through brain or tumor cell membranes that the drug has to cross to reach the target site [23]. Numerous studies have evaluated the composition of the neurons and brain cells considering age or the regional distribution in different domains, as the main factors [24–27]. Human brain is composed of specific cells with membranes formed by several layers, each of different lipid components [28]. TMZ has been used for the treatment of glioblastoma which is a tumor most commonly located in the supratentorial region of the brain [29]. However, up to the best of our knowledge, no studies concerning the structural interactions of TMZ with biological brain plasma membranes have been reported yet. In this work, we have considered a realistically complex lipid model of a human neuronal/brain plasma membrane (BPM) which can represent with good accuracy the main components of the real system. A general scheme of the membranes forming the external layers of brain cells is sketched in Figure 1. We should note that the inner and outer leaflets of BPM have different lipid compositions.

**Fig 1.**
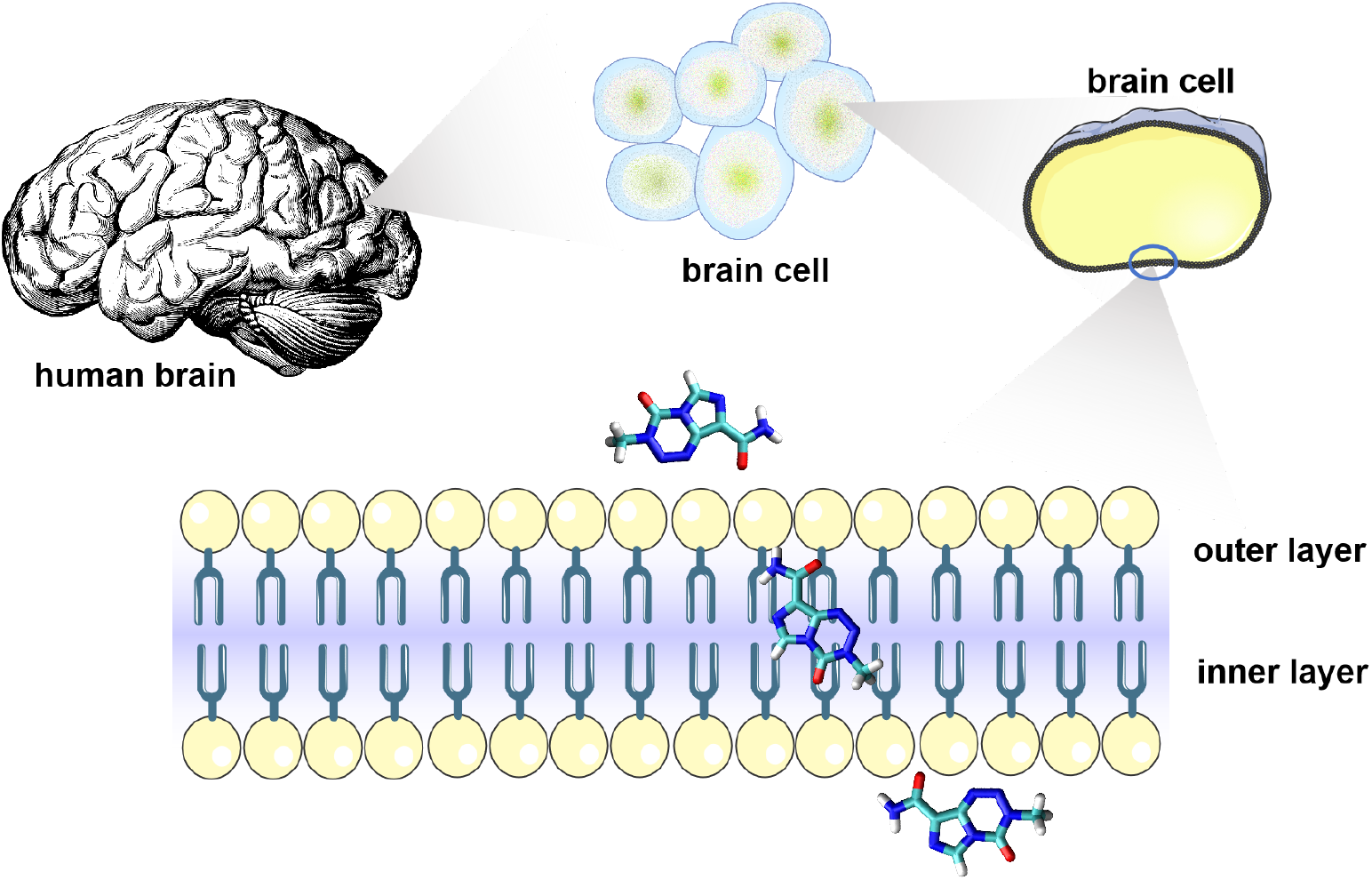
TMZ crossing brain cell membranes. Schematic diagram of the process of TMZ crossing the BPM. The brain bilayer cell membrane contains a wide variety of lipids, with the types and proportions of lipids in the upper and lower layers showing some significant differences.

In a pioneering work in computational modeling of the brain plasma membrane, Ingólfsson et al. [30] constructed a coarse-grain (CG) model for the BPM composed of tens of lipids of different classes, based on previous knowledge from a number of studies of lipidomic measurements of neurons and brain tissue. In the present work, we have considered the main types of lipids reported in Ingólfsson et al.’s work [30] and selected the most abundant and representative one from each group class. This has allowed us to perform simulations at the all-atom level without the need of considering a number of atoms too large and unpractical. In this work, we have focused our efforts on the study of the free energy barriers involved in the translocation of TMZ through the external layer of the BPM. Once the drug has crossed this barrier, chemical processes occur and TMZ follows the chemical activation described above. In summary, the present study is devoted to the analysis of the preferential sites and orientations of TMZ at the aqueous solution (representing blood plasma), and at the interface of the brain cell membrane. Nevertheless, the main result of this paper is the calculation of the free energy barriers that TMZ should pass through the BPM as modeled in a realistic way. The work is organized as follows: methods and computational details are described with all details in Section; the main results and the corresponding discussion are described in Section and the main conclusions are reported in Section. Finally, a few relevant complementary properties are reported in the ”Supporting Information” (SI).

## Methods

We obtained the structure of TMZ (PubChem CID: 5394) from the National Center for Biotechnology Information PubChem database (http://www.ncbi.nlm.nih.gov/pccompound) [9]. A realistic model of the brain cell membrane has been generated with the well-known CHARMM-GUI web-based tool [31, 32]. Sketches of the backbone structures of TMZ and the main components of the brain membrane are represented in Fig 2. The model membrane used in this work is a simplified but reliable representation of the mammalian brain plasma membrane, comprising two leaflets (outer and inner layers) with different compositions. A pioneering computational modeling based on CG simulations [30] found significant differences between the normal cell and brain cell membranes, the latter formed by a mixture of diverse lipids, sterols and other organic substances: phosphatidylcholine (PC) lipids, phosphatidylethanolamine (PE) lipids, sphingomyelin (SM) lipids, phosphatidylserine (PS) lipids, glycolipids (GM), cerebrosides, phosphatidylinositol (PI) lipids, phosphatic acids (PA), phosphatidylinositol phosphates (PIP), ceramides (CER), lysophosphatidylcholine (LPC) lipids, lysophosphatidylethanolamine (LPE) lipids and the sterols ciacylglycerol (DAG) and cholesterol (CHOL). We considered Ingólfsson et al.’s [30] model to analyze the interactions of lipids with TMZ. To simplify the study, we selected the most abundant lipid to represent one lipid type, neglecting those with minimal percentages. Subsequently, we constructed a 400-lipid membrane based on the percentages indicated in Table 1. It is important to note that we employed all-atom molecular dynamics (MD) simulations, providing a higher accuracy compared to CG, due to the atomic-level description.

**Table 1.** Composition of the two leaflets (outer, inner) of the brain membrane, with percentages and number of lipids *N*.

| Class | Species | Outer layer (%) | $N_{outer}$ | Inner layer (%) | $N_{inner}$ |
| --- | --- | --- | --- | --- | --- |
| PC | DPPC | 24.2 | 48 | 13.6 | 27 |
| PE | POPE | 11.0 | 22 | 21.3 | 43 |
| SM | DPSM | 11.0 | 22 | 6.0 | 12 |
| **PS** | **POPS** | **0.0** | **0** | **14.5** | **29** |
| **cerebrosides** | **DPGS** | **9.4** | **19** | **0.0** | **0** |
| CHOL | CHOL | 44.4 | 89 | 44.6 | 89 |
| Total | - | 100.0 | 200 | 100.0 | 200 |

**Fig 2.**
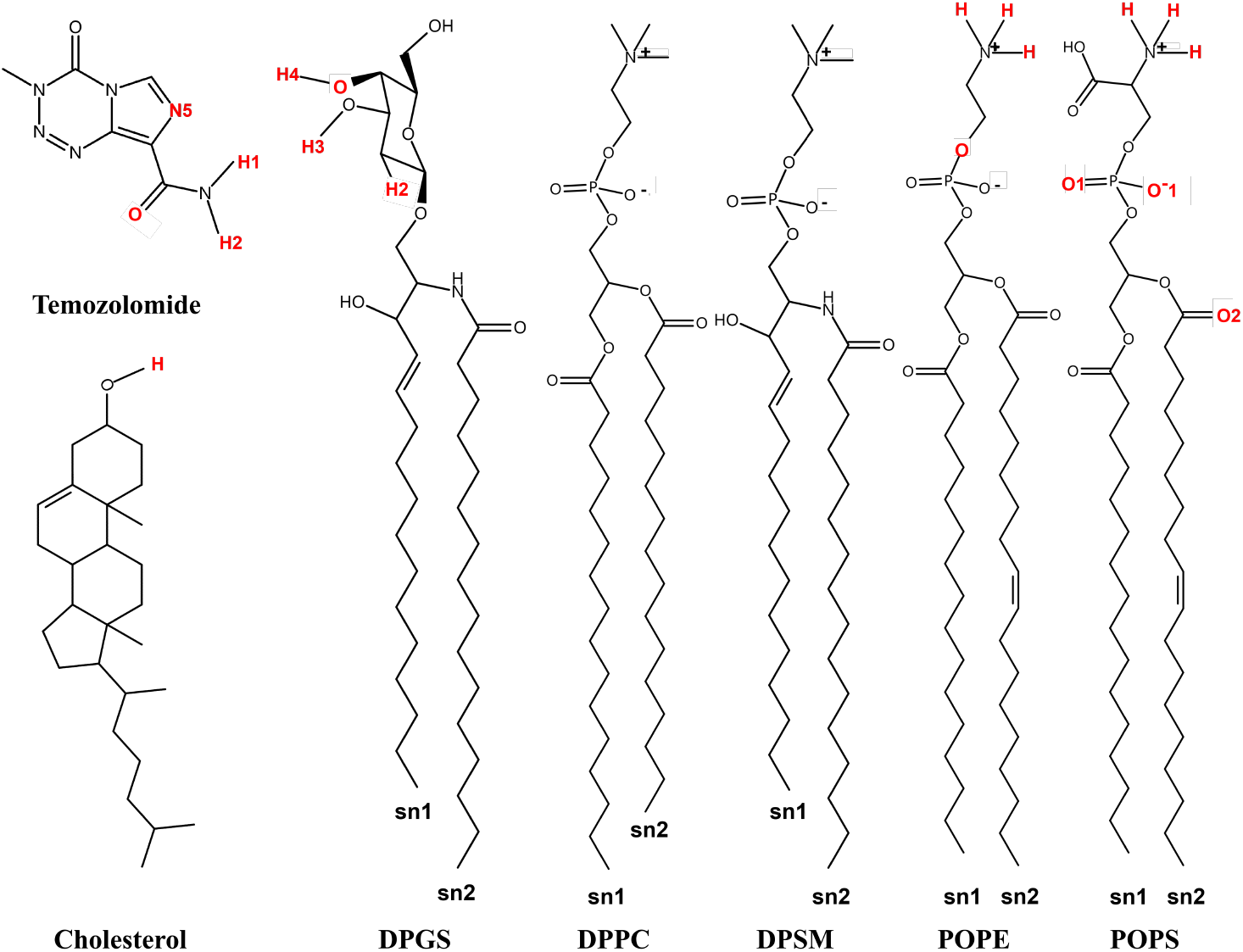
Backbone structures. Sketches of molecular structures of TMZ and the main classes of lipids considered in this work: cholesterol (CHOL), di-palmytoil-phosphatidylcholine (DPPC), 1-palmitoyl-2-oleoyl-sn-glycero-3-phosphoethanolamine (POPE), 1-palmitoyl-2-oleoyl-sn-glycero-3-phospho-L-serine (POPS), N-(hexadecanoyl)-hexadecasphing-4-enine-1-phosphocholine (DPSM) and 1,2-dipalmitoyl-sn-glycero-3-succinate (DPGS). The latter are cerebrosides and they are lipids exclusively located at the brain cells. Part of hydrogen-carbon bonds are not shown, for the sake of clarity. The highlighted sites will be referred to in the text by the same labels.

The NAMD2 software package [33] with the reparameterized CHARMM36m force field [34–37] was used in all MD simulations at two temperatures of 310.15 K and 323.15 K. TMZ and lipids are fully solvated by 53,556 TIP3P water molecules and sodium chloride at 0.15 M concentration, yielding a system of 95,235 atoms. The initial system size is 101 Å*×* 101 Å*×* 100 Å, and periodic boundary conditions are all applied in *X, Y*, and *Z* directions. The *XY* plane is taken along the surface of the BPM, whereas *Z* is the direction normal to the instantaneous *XY* plane. The Van der Waals interactions included a smoothing function starting at 10 Å and its cutoff was 12 Å, with a pairlist distance of 16 Å. Long-range electrostatic forces were computed using particle mesh Ewald [38], with a grid space of 1 Å. Periodic boundary conditions in all spatial directions are considered. A Langevin thermostat [39] was used with a damping coefficient of 1 ps^−1^ to control the temperature, and the pressure was set at 1 atm and regulated by a Nose-Hoover Langevin piston [40] with Langevin dynamics [41] at an oscillation period of 50 fs. After 100 ns equilibration periods at the NVT ensemble, 0.5 *μ*s trajectories with a timestep of 2 fs were generated at the NPT ensemble for each system. All bonds involving hydrogens were set to fixed lengths, allowing fluctuations of bond distances and angles for the remaining atoms.

## Results and Discussion

### Physical characteristics of BPM and TMZ at its interface

To describe the phase states of the BPM simulated in the present work, we have computed the so-called deuterium order parameter *S*_*CD*_, as it was defined in references [42–45], which can help us to efficiently characterize the ordering inside the hydrated lipid bilayer. *S*_*CD*_ is defined for each *CH*_2_ group of the two tails of each lipid class as follows:

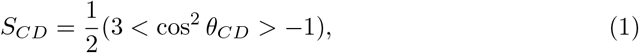

where *θ*_*CD*_ is the angle between the direction normal to the surface of the membrane and a CH-bond. It is remarkable that *S*_*CD*_ can be obtained from ^2^H NMR experiments [46], making it a suitable property to verify the reliability of the simulations. The averaged results are shown in Fig 3 for both tail chains of several classes of lipids considered in this work. The results indicate that all classes of lipids investigated show maxima of *S*_*CD*_ between 0.37 and 0.43 units, namely with lower values found at 323 K and higher at 310 K. These values larger than 0.4 are in overall good agreement with the results of Ingólfsson et al. [30] for the full brain cell CG model reported by such authors. Since *S*_*CD*_ is an indication of the degree of ordering (lineality) of the acyl chains, we confirm that the acyl chains of all types of lipids exhibit slightly greater ordering at 310 K, which becomes more disordered at 323 K, as expected. On the other hand, we can observe two additional features: (1) the ordering at the inner layer (triangles) is slightly greater than at the outer layer (circles) which indicates a more rigid phase for the inner leaflet and (2) those lipids having unsaturated bonds in one of their acyl tails, such as POPE, DPSM, POPS, and DPSG show highly disordered profiles of *S*_*CD*_ at the double bond positions.

**Fig 3.**
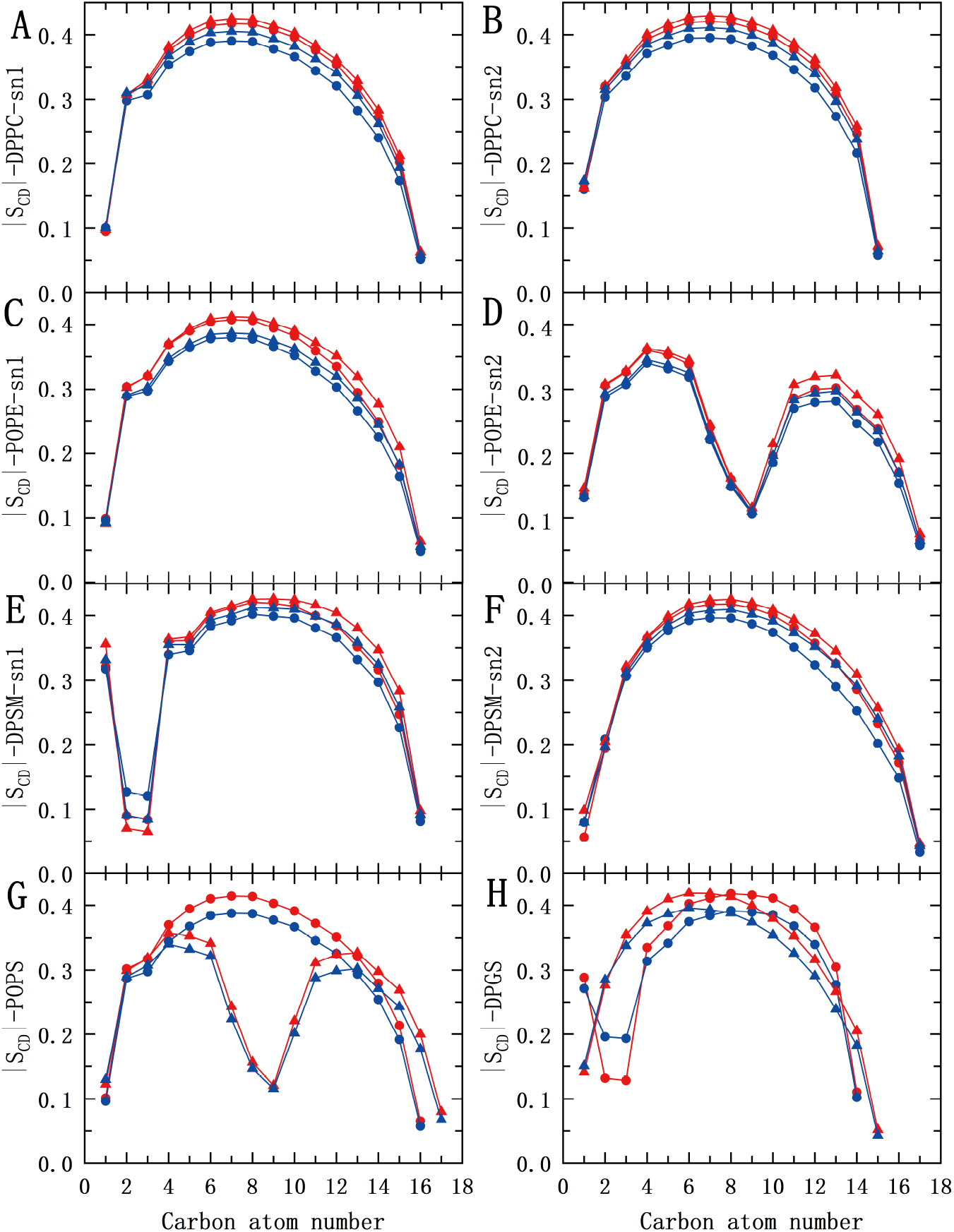
Order parameter. |*S*_*CD*_| for two acyl tails (sn1 and sn2 shown in Fig2) of selected lipid species considered in the membrane model of brain cells: DPPC (A-B); POPE (C-D); DPSM (E-F); POPS (G) and DPGS (H). Lipids at the inner layer are indicated with triangles and those at the outer layer are represented with circles. POPS in panel G is present only in the inner layer, while DPGS in panel H is only in the outer layer, and the circle represents sn1 while the triangle represents sn2. Systems are also represented at temperatures of 310 K (red) and 323 K (blue).

Other relevant parameters for the control of the simulations are the area per lipid *A* and thickness Δ*z* of the membrane. *A* was obtained considering the membrane surface along the *XY* plane divided by the number of lipids and cholesterol [47]. The two leaflets contain the same number of lipids and their *XY* plane surfaces are the same at each instant of the simulation. Averaged area per lipid of the two membranes (low, high temperature) are reported in Table 2. We obtained values of *A* around 41 Å^2^ at 310 K and slightly higher at 323 K, indicating a larger thermal disorder at the latter case. The full profile of *A* as a function of the simulation time is reported in Section ”Supporting Information”, Fig. S1. Our results are in good overall agreement with other computational works, such as the reference work of Ingólfsson et al. [30] or the study of Yee et al. [48] on the BPM of a healthy brain compared to another with Alzeheimer’s disease. The values of *A* by these authors were of 46 and 42.3 Å^2^ respectively, i.e. with discrepancies between 3 and 11%, probably due to the different models considered.

**Table 2.** Area per lipid (in (Å^2^) and thickness (Δ*z* (in Å))of the membrane at 310 and 323 K. *A* and Δ*z*. Estimated errors in parenthesis.

| Property | 310 K system | 323 K system |
| --- | --- | --- |
| Thickness ($\text{\AA}$) | 42.63 (0.19) | 42.13 (0.23) |
| Area per lipid ($\text{\AA}^2$) | 41.24 (0.24) | 42.44 (0.32) |

The thickness of the membrane may provide additional clues about the influence of cholesterol on the mechanical properties of plasma membranes, such as rigidity and capability of allowing the movement of species in and out of the cell. We have obtained the thickness of the membrane Δ*z* as defined in Fig. S2 of SI. The results reported here (around 42 Å at 310 K) are again qualitatively close to those reported in Refs. [30] (41 Å) and Ref. [48] (47.3 Å) where more detailed models were considered, but only reported at 310 K. The full profile of Δ*z* as a function of the simulation time is reported in Section ”Supporting Information”, Fig. S2 showing fluctuations of the order of 5% around the averaged values, reported in Table 2, where we can observe that as the temperature increases from 310 K to 323 K, thickness slightly decreases, while area per lipid increases. These results indicate a slightly higher degree of disorder as temperature increases, given the higher values of *A* and smaller values of Δ*z* at 323 K as expected.

In passing, we should mention that detailed studies on the orientational order of TMZ at the BPM’s interface indicate a strong tendency of TMZ to stay oriented in a particular fashion, i.e. with the dihedral angle *θ* formed by atoms N5-C4-C5-O1 of TMZ taking values around cos *θ* ∼ 0.9. For full details see Fig. S3 in SI. As an additional property that can be relevant for the understanding of the behavior of TMZ at the interface of the BPM, we have evaluated the Z-axis position of TMZ along the simulations at normal body and high temperatures and observed a marked tendency of TMZ to stay at the interface and, more preferentially, solvated in the aqueous ionic solution, with no incursions in the central region of the membrane. To illustrate this fact, we report the Z-axis position of TMZ as a function of the simulation time in Fig. S4 of SI.

### Radial distribution functions of TMZ bound to cholesterol and selected lipids

A direct route to the characterization of the local structure of each atomic species of the system is usually obtained by means of normalized radial distribution functions (RDF) *g*_*AB*_(*r*) for two different species *A* and *B* (see Eq.(2) of Ref. [45], for instance), indicating the probability of finding a species *A* at a given distance *r* of a species *B*. Among the wide variety of possible RDF that could be computed, we have considered only the most relevant RDF based on the first coordination shells of TMZ with cholesterol and selected lipids. The remaining RDF indicate low maxima at distances significantly longer than the typical hydrogen bond (HB) values or show too noisy profiles which indicate that the corresponding local structures are not stable enough.

As a general fact, we can observe a neat first coordination shell with a sharp maximum located around 1.8-2.0 Å that is the signature of HB between TMZ and the remaining species. This range of distances is typical of the HB connections between small-molecules and organic species (see Ref. [49] and references therein). Among all lipids, the first peak in the radial distribution function of POPE/POPS with N5 of TMZ is significantly higher than others, especially for hydrogens located at the headgroups of POPE/POPS (see Fig. 4, panels C and D), indicating strongest bindings of TMZ to phosphatidylethanolamine and phosphatidylserine lipids. Conversely, panels B and E in Fig. 4 show that DPGS, the special cerebroside typical of the outer leaflet of brain cell membrane, also contributes to the binding of TMZ to the membrane through hydrogens H3 and H4, relatively stronger interaction between TMZ and DPGS is shown at the temperature of 323 K. The noisy profiles of panels B and E indicate that the lifetime of the TMZ-DPGS interaction is of shorter duration than in TMZ-POPE/POPS binding, because the long-lived interactions (namely HB) produce smoother RDF profiles than short-lived ones, which suffer abundant breaking and reforming events.

**Fig 4.**
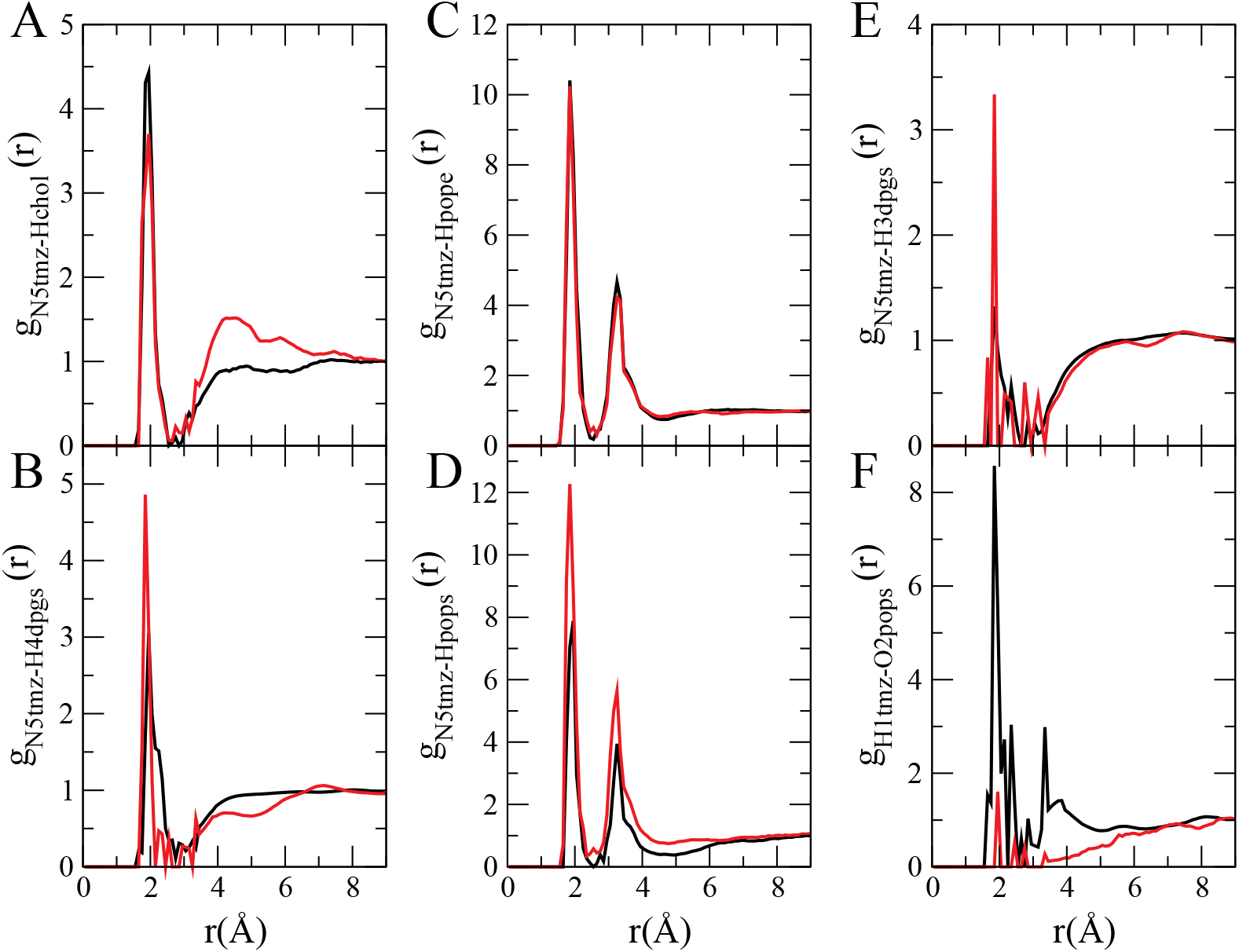
Radial distribution functions. RDF for different atoms in TMZ with several lipids at 310 K (black) and 323 K (red) systems. In panel A, *g*_*N*5*tmz*−*Hchol*(*r*)_ refers to the RDF of the atom named N5 (see Fig. 2) in TMZ and the hydrogen of the hydroxyl group of cholesterol. In panels B-F the selected atoms are also indicated with the same labels as in Fig. 2.

In the particular case of TMZ-DPGS hydrogen-bonding we can observe mainly HB of nitrogen N5 of TMZ with H3 and H4 from DPGS and a oxygen-hydrogen interaction of H1 of TMZ with O2 in POPS, but only well defined at 310 K. Comparing systems at 310 K and 323 K, it can be noticed that temperature plays a minor role in most cases. In a few cases, such as TMZ-DGPS bindings (see panels B and E of Fig. 4) we can notice an enhancement of the first maxima at 323 K, that can be attributed to more frequent and durable events of TMZ-DPGS pairings than at 310 K, when TMZ shows a bigger affinity to bind cholesterol (panel A in Fig. 4) and POPS lipids (panel F of Fig. 4).

### Estimation of free energy barriers for TMZ crossing

From a general perspective, the calculation of the Helmholtz or Gibbs free energy differences for binding processes or for configurational changes is a difficult task and it requires a considerable amount of computer time and a precise knowledge of the hypersurface of potential energy of the system [50]. This can be explored by means of methods such as metadynamics [51–53], hybrid quantum mechanics/molecular mechanics methods [54] or transition path sampling [55–58]. However, a usual way to obtain free energy estimations is through the calculation of potentials of mean force (PMF) through a variety of methods such as umbrella sampling [59], constrained molecular dynamics [60] or adaptive biasing force methods, among many others.

We calculated free energy profiles by the adaptive biasing force (ABF) method [61, 62], where the PMF is derived from integrating the mean force F_*z*_ over the reaction coordinates *ξ*_0_ [63–65]. The total work through the membrane is:

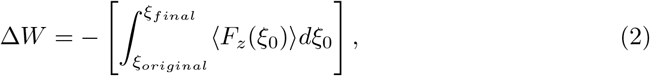

where *ξ*_*original*_ is the original reaction coordinate of the TMZ and *ξ*_*final*_ is the final reaction coordinate of the TMZ. *(F*_*z*_(*ξ*_0_)) is the average force in the z-direction exerted on the TMZ at *ξ*_0_. The ABF method offers the advantage of not requiring the estimation of free energy barriers and providing more diverse sampling. In typical simulations, the substrate is driven in a specific direction by the total force acting on it. With ABF, a biasing force in the opposite direction is introduced, creating a balance of the substrate and leading to random moving. The reaction coordinates in this case represent the distance along the z-axis from the center of mass of TMZ to a ”dummy atom” located at the point (0.0, 0.0, 0.0). These collective variables in small size improve the performance of ABF calculation. The reaction coordinates are divided into several windows. TMZ will be confined within specific windows by harmonic forces in the Z-axis direction at the boundaries, which enhances the sampling efficiency. Two TMZ membrane-penetrating ABF systems were set up at 310 K and 323 K respectively. Each window has a width of 5 Å, and the PMF is calculated every 0.1 Å. The force constant at the window boundaries is of 20 kcal/mol/Å^2^. A sample count of the times that TMZ has been placed at different positions in the system is reported in Fig. S5 of SI to proof the reliability of the ABF calculations.

To obtain the free energy profile of TMZ for the entire membrane, the process is divided into two sections: (1) from the extramembrane to the center of the bilayer (Process A) and (2) from the intramembrane to the center of the bilayer (Process B) in order to prevent possible artificial effects. To clarify the PMF curves of A and B along the reaction coordinate, the Y-axis of the A curve is shifted so that the PMF value at the initial reaction coordinate (aqueous solution outside the membrane) is 0. The A and B curves were combined at the center of the membrane, and the Y-axis of curve B was shifted to align the PMF values of A and B at the bilayer center. In other words, we converted the reaction coordinates into the corresponding difference in location along he Z-axis between the center of mass of TMZ and the center of the membrane. (named as ”Z-distance”). For the sake of clarity, we identified the center of mass of cholesterol in the bilayer as the center of the bilayer. Our results are shown in Fig. 5. The PMF profile of the system at 310 K indicates a local energy minimum occurring at when the Z-distance is approximately -4 Å. To verify this, we conducted unbiased MD simulations at the NPT ensemble, initially placing TMZ around the center of the membrane and the local energy minimum position. The temperature was set to 310 K with a small timestep of 0.5 fs, and each simulation was repeated twice. With this procedure we explored the likeliness of TMZ to move in or out the membrane as a function of its initial location.

**Fig 5.**
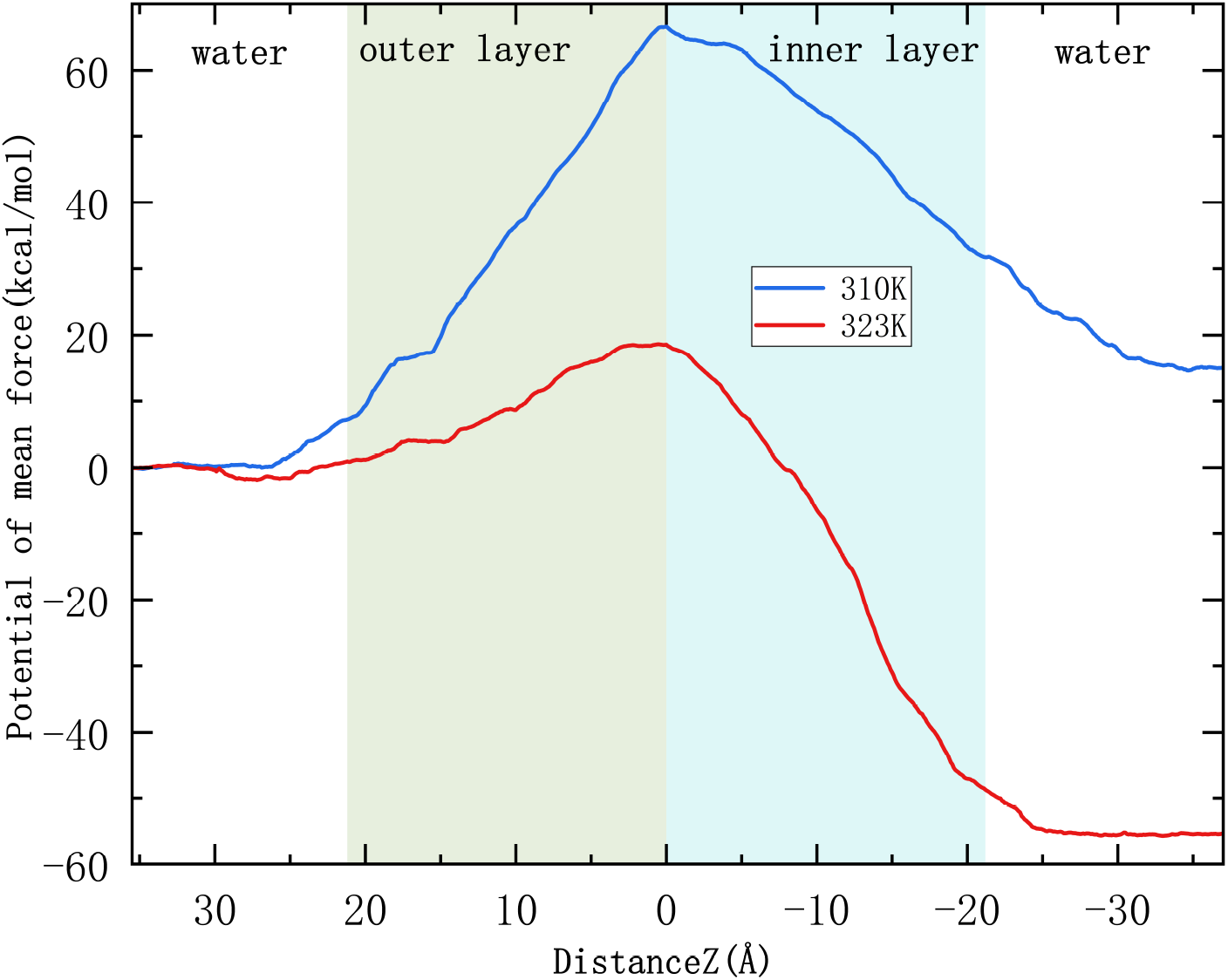
Free energy profiles of TMZ crossing the brain cell membrane bilayer. The origin of the free energy (potential of mean force) is taken when TMZ being solvated in the extracellular matrix, namely the aqueous solution. For each system, we calculated the PMF of TMZ from the extracellular matrix to the center of the membrane and the PMF of TMZ from the intracellular matrix to the center of the membrane, and merged the two curves at the center of the membrane (*Z* =0 Å;), where the energy is highest, in order to set a common reference point.

The energy barrier for TMZ to pass through the upper layer at 323 K is significantly lower than that at 310 K. On the contrary, the energy released when TMZ passes through the lower layer in the 323 K system is larger than that of 310 K. In the 310 K system, the free energy has a local minimum when TMZ is in the inner leaflet of the BPM and locates around 0 Å. A way to rationalize these findings can be described as follows.

We have started four independent unbiased simulations where TMZ started around the center of the membrane at 310 K, with the corresponding initial configurations of these simulations taken from trajectories of ABF calculations. The results for the TMZ position as a function of time are shown in Fig. 6, where each trajectory has been labeled as ”trajectory 1-4”. We observed that when the original position of TMZ is just at the center of the lipid around 0 Å, TMZ tends to cross the outer layer to the water layer, and the process is within 20 ns; when TMZ is placed at around -4 Å, TMZ tends to cross the inner layer, and the process takes around 40 ∼ 50 ns. The free energy calculation results show that at 310 K, the energy released by TMZ in the center of the lipid is higher when it passes through the upper leaflets. This might be be the reason why TMZ at 0 Å tends to pass through the upper leaflets and at a faster speed. On the contrary, the energy difference of TMZ through the lower leaflets is smaller, and the transmembrane process is slower. When the original position of TMZ is around -4 Å, TMZ first moves to the outer layer to a position of about 5 Å, and then passes through the inner layer. The energy calculation results of the 310 K system show that a local energy minimum appears around ∼ -3 Å, which may be the reason why TMZ moves upward first.

**Fig 6.**
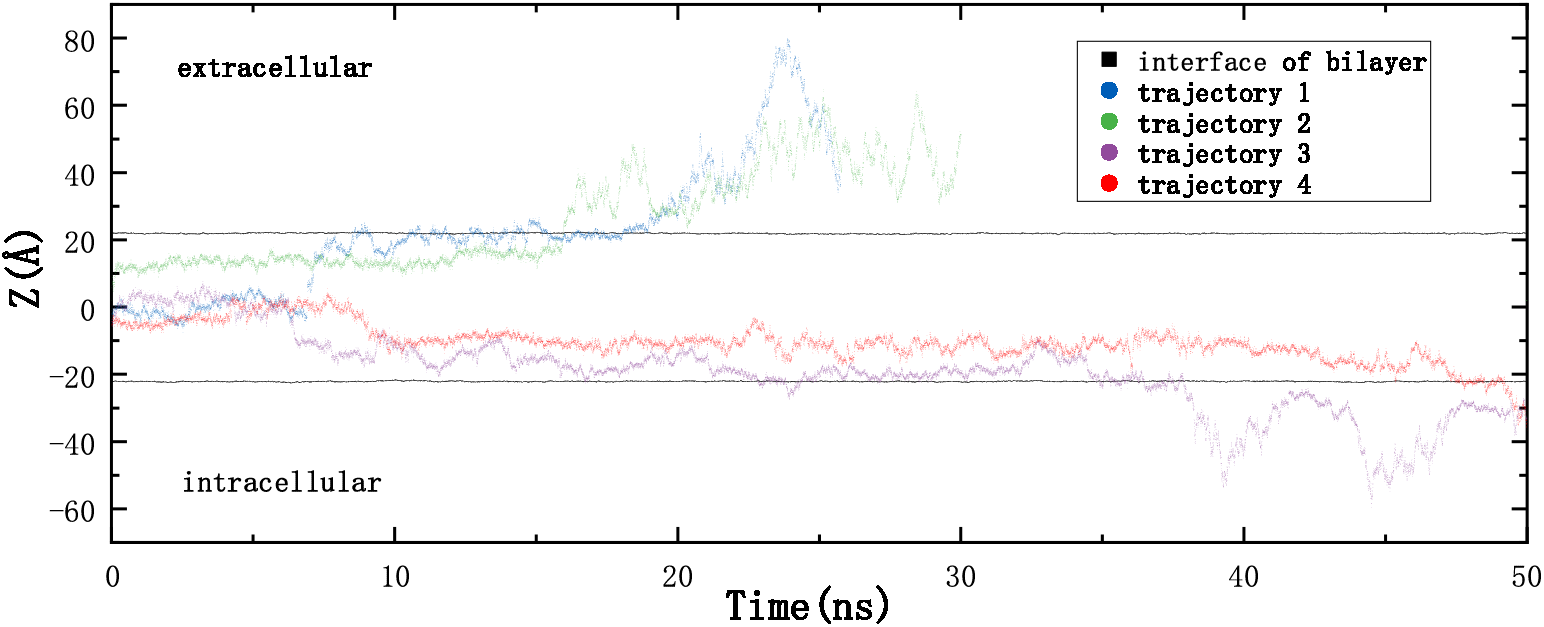
Z-axis position of TMZ. Displacement of TMZ inside lipid bilayer for four different trajectories. Circles indicate the position of the center of mass of TMZ as defined in Fig. S2 of SI. The initial location of TMZ inside the membrane has been set between (0 and -4 Å). Black lines stand for the average position of phosphorus atoms of DPPC, DPSM, POPE, POPS, oxygen atoms of cholesterol, and O4 of DPGS in each layer.

Finally, in order to find additional structural clues to the striking decrease in the free energy barrier from 65 to 20 kcal/mol after a moderate rise of temperature of 13 degrees, we report additional RDF of TMZ with cholesterol, DPGS, POPS and POPE in Fig. S6 of SI. Interestingly, we have obtained these RDF during the computation of free energies, i.e. from configurations associated to the ABF calculations, in order to find out the preferential binding of TMZ when crossing the BPM. We should remark that these RDF are different of the RDF reported in Section, obtained from unbiased MD configurations. From the RDFs reported in Fig. S6 (A) and (B), we have found at 310 K evidence of strong pairings of TMZ with cholesterol and, in a weaker fashion, to DPGS (see Fig. S6 C), whereas the interactions with POPE are much weaker and no signature of TMZ-POPS is observed. Differently, at 323 K TMZ shows a tendency to be mainly bound to POPE (see Fig. S6 D) but also to POPS and DPGS and no traces of bindings to cholesterol. The latter fact is quite surprising and it can be due to the possible formation of cholesterol clusters at 323 K as indicated in Ingólfsson et al.’s work. All in all, the main reason of the strong reduction of the free energy barrier should be attributed to the strong TMZ-cholesterol interactions at the normal human body temperature. After this observation, we can suggest that cholesterol and DPGS are the main controlling agents for the permeation of TMZ. Consequently, as a rough indication, a patient suffering brain tumor might be bound to smaller intake of food containing large amounts of cholesterol or cerebroside-like lipids (such as DPGS) [66] to reach faster TMZ crossing of the BPM. In other words, limiting the intake of the above mentioned compounds would be a primary action allowing a patient to use smaller doses of TMZ in order to get a similar medical effect and, additionally, to achieve a reduction of the well-known side effects caused by TMZ, as described in Ref. [17].

## Conclusions

In the present study, a comprehensive computational analysis characterizing the permeation of TMZ across the brain-like plasma membrane bilayer was conducted. We have introduced a TMZ drug molecule at the interface of a model BPM and systematically observed its interactions with solvent water, ions, and the distinct regions of the membrane, which comprise different outer and inner layers. Our results confirm the correct equilibration of the system and structural data indicate that the characteristic order parameter used to verify that the system is in a liquid ordered state and other structural properties such as the area per lipid and thickness of the membrane are in fair qualitative agreement with the results reported by Ingólfsson et al. [30] and by Yee et al. [48], where somewhat more sophisticated models of the BPM were used.

Our data of the free energy profiles for TMZ transmembrane crossing obtained from adaptive biasing force methods, indicate the highest affinity of TMZ to stay at the interface of the BPM with the aqueous solution, instead of spontaneously cross the membrane by pure diffusion. This indicates that in the real system, some agents such as membrane ABC tranporters [67] or SLC transporters [68, 69] may play a key role for the crossing of TMZ through the external layer of the brain cell membrane. Interestingly, when TMZ has reached the internal hydrophobic regions of the BPM, small fluctuations in its position can lead to a successful crossing to the internal regions and to eventually reach the first cytoplasmic region of the brain. Our ABF calculations indicate that TMZ needs about 65 kcal/mol to cross the BPM at the normal human body temperature of 310 K, whereas this barrier decreases to about 20 kcal/mol at 323 K. Data obtained by RDF indicate that the formation of strong TMZ-cholesterol and TMZ-DPGS pairings at 310 K would make it more difficult for TMZ the permeation through the BPM barrier. In addition, those preferential interactions of TMZ with cholesterol and DPGS over the other classes of lipids, would indicate the potentially positive effect of reducing the intake of food with high contents of such species in glioblastoma patients. This would allow them an easier permeation of TMZ through the BPM eventually reducing the dose and, consequently, the associated side effects.

## Supporting information

Supplementary Materials for the article

## Acknowledgments

The authors acknowledge financial support provided by the Spanish Ministry of Science, Innovation and Universities. Huixia Lu thanks the financial support by the ”Margarita Salas” grant which is funded by the European Union – NextGenerationEU. This publication is a part of the I+D+i project with reference PID2021-124297NB-C32, founded by MCIN/AEI/10.13-039/501100011033 and “FEDER Una manera de hacer Europa”. Yanhong Ge is a Ph.D. fellow from the China Scholarship Council (grant 202306230043). J.M. thanks the *Generalitat de Catalunya* for the support through the grant *GrupdeRecercaSGR* − *Cat2021Condensed, ComplexandQuantumMatterGroup* reference 2021SGR-01411 and to the Polytechnic University of Catalonia-Barcelona Tech through the funding AGRUPS. The authors thankfully acknowledges the computer resources at MareNostrum5, Storage5 and the technical support provided by BSC (RES-BCV-2024-2-0006).

## Data availability statement

All related files including an example used to calculate the binding free energy of this work can be found in our repository of Zenodo: 10.5281/zenodo.12745251.

