## Supplementary Materials for the article for "Free energy profiles of temozolomide crossing brain plasma membranes"

### Supporting Information

In this section we report detailed results of the area per lipid and thickness of the model brain plasma membrane (BPM) and the angular orientations and Z-axis positions of TMZ, all as a function of time at the two temperatures considered in the work (310 and 323 K). Finally, a counting test of the adaptive biasing force (ABF) method and specific radial distribution functions (RDF) of TMZ with lipids are also reported.

#### Area per lipid and thickness of the BPM

The area per lipid of the BPM as a function of time is reported in Fig. S1 and the thickness of the BPM is shown in Fig. S2.

#### Angular orientations and positions of TMZ at the surface of the BPM

In order to analyze the angular distributions of TMZ at the interface of the membrane, we have defined three sorts of angles, totally equivalent, so that we chose only one of them for analysis. In all cases, we can observe that the orientation of the dihedral angle $\theta$ formed by atoms N5-C4-C5-O1 of TMZ is nearly constant at all times at around $\cos\theta \sim 0.9$ and it only suffers sudden fluctuations of the dihedral angle to values close to $\cos\theta \sim -0.3$, as indicated in Fig. S3. We can attribute those flips to the contacts between TMZ and the lipids and the interface of the membrane.

The penetration of TMZ in the membrane along its normal direction is a very relevant feature. We report in Fig. S4 the Z-axis position of TMZ from the center of the bilayer (i.e. $z = 0$) using the last meaningful 200 ns of each production trajectory. It stands clearly that TMZ stays at the aqueous solution surrounding the membrane most of the time, with a few occasional visits to the interface of the membrane.

#### Test of the adaptive biasing force method

A detailed count of the ABF method in order to check its suitability is reported in Fig. S5.

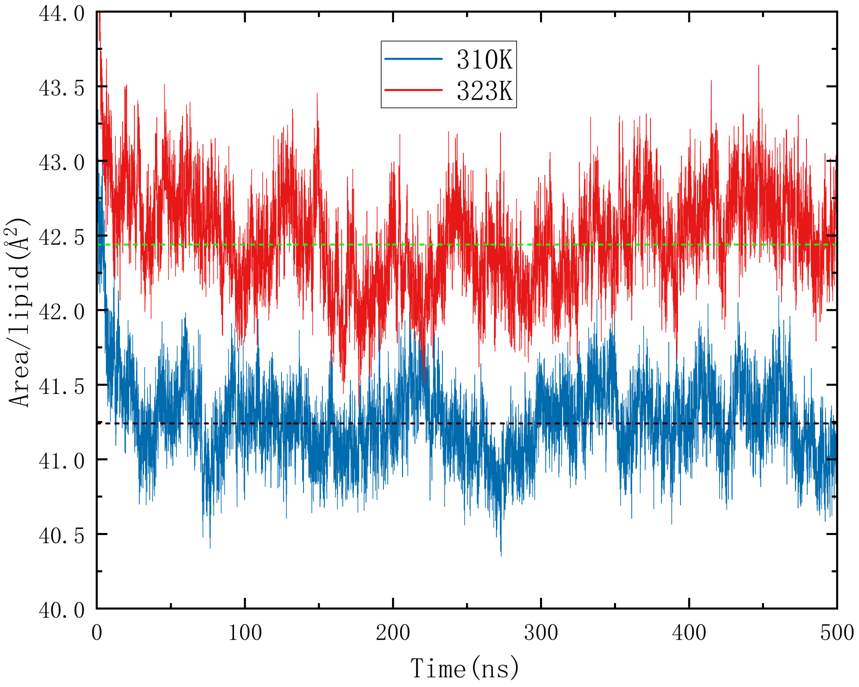


**Fig S1. Area per lipid.** The area per lipid of the brain membrane as a function of simulation time in the 310 K and 323 K systems. Dashed lines correspond to their average values in the last 400 ns.

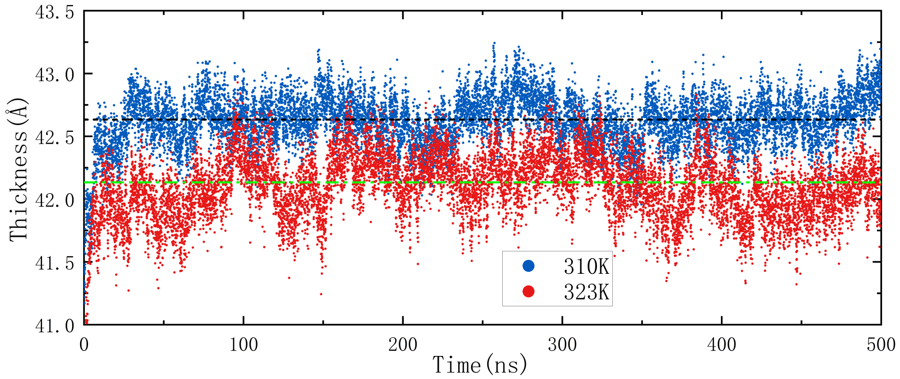


**Fig S2. Thickness.** The thickness of the brain membrane as a function of simulation time in the 310 K and 323 K systems. Dashed lines correspond to their average values in the last 400 ns.

### Additional radial distribution functions of TMZ-lipids

In Fig. S6 we report additional RDF of TMZ with cholesterol, DPGS, POPS and POPE. We should indicate that these RDF have been computed from configurations associated to the ABF calculations and are different of the RDF obtained from unbiased MD configurations in Fig. 4 of the main text.

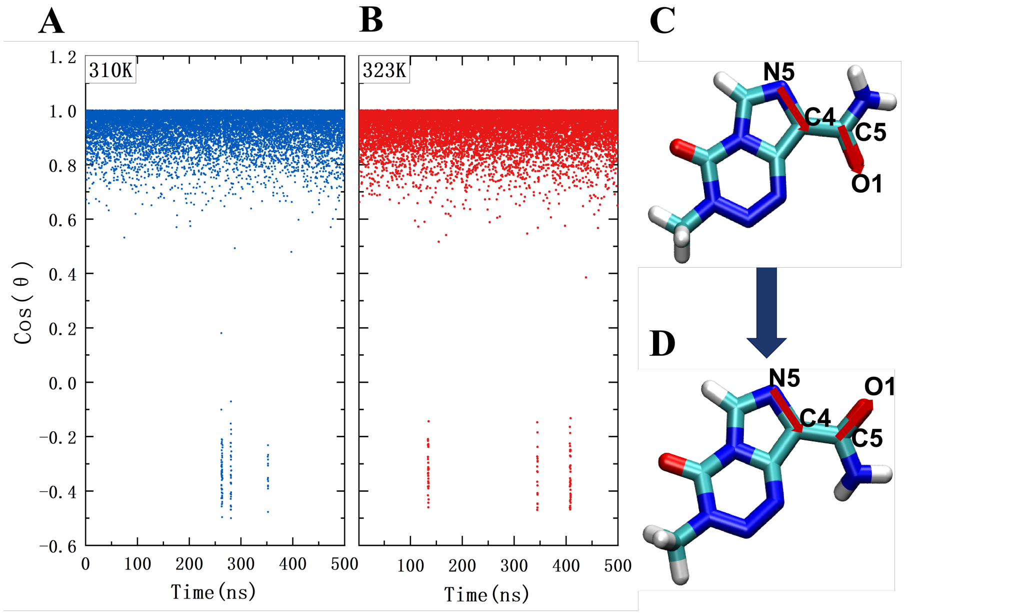


**Fig S3. TMZ occasionally flips along the simulation.** The dihedral (torsional) angle $\theta$ is defined by the sites N5-C4-C5-O1 indicated in C by the two red arrows. Panels A and B show the dihedral evolution along the simulation time for two systems studied in this work; Panel C shows two different configurations of TMZ: the upper subfigure presents a stable structure of TMZ with Cos$\theta$ $\sim$ 0.9 and the lower subfigure refers to a flipped structure of TMZ with Cos$\theta$ $\sim$ -0.3.

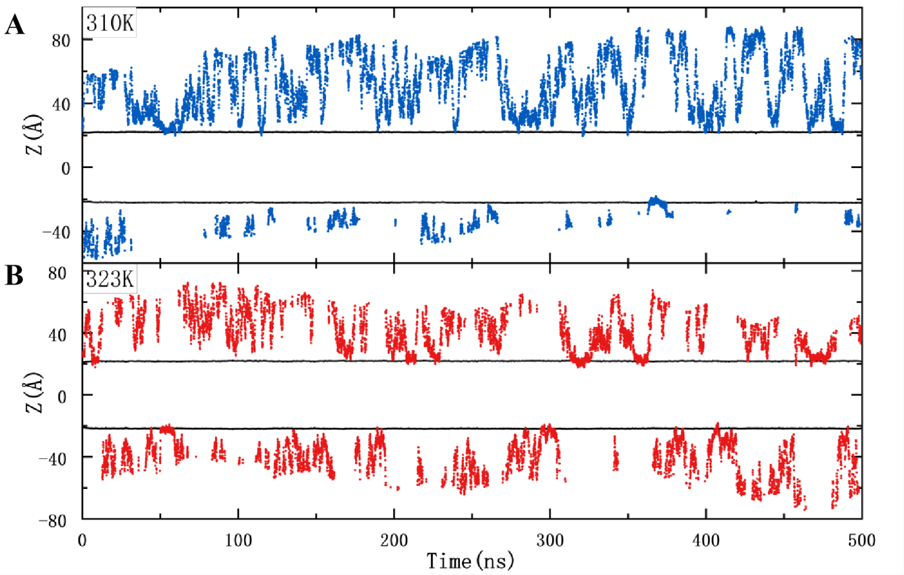


**Fig S4. Z-axis position of TMZ in 310 K and 323 K systems.** The same definition of the interface of membrane is utilized here as in Fig. 6 of the main text. Penetration of TMZ inside the external brain bilayer (circles indicate the position of the center of mass of TMZ whereas squares stand for the average position of phosphorus atoms in DPPC, DPSM, POPE, POPS and O3 in cholesterol and O4 of DPGS in each layer.

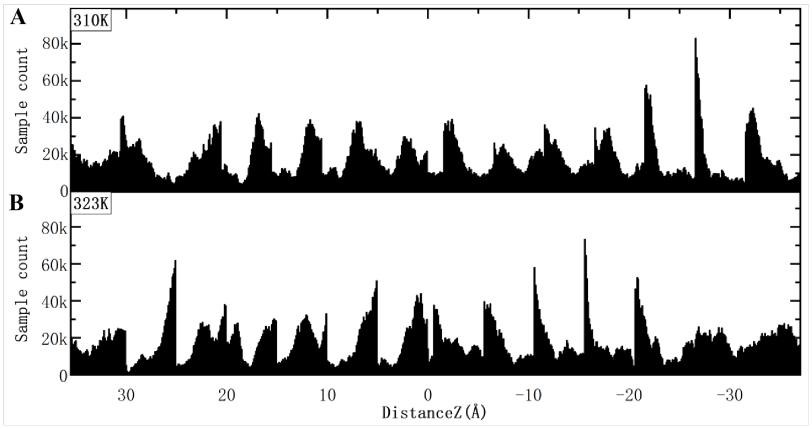


**Fig S5. Sample count of potential of mean force in 310 K and 323 K systems.** "Distance Z" refers to the distance on Z-axis between the center of weight mass of TMZ and a dummy atom (0, 0, 0), refering to the membrane center. The sample count is always above 1000, which makes the calculation result reliable.

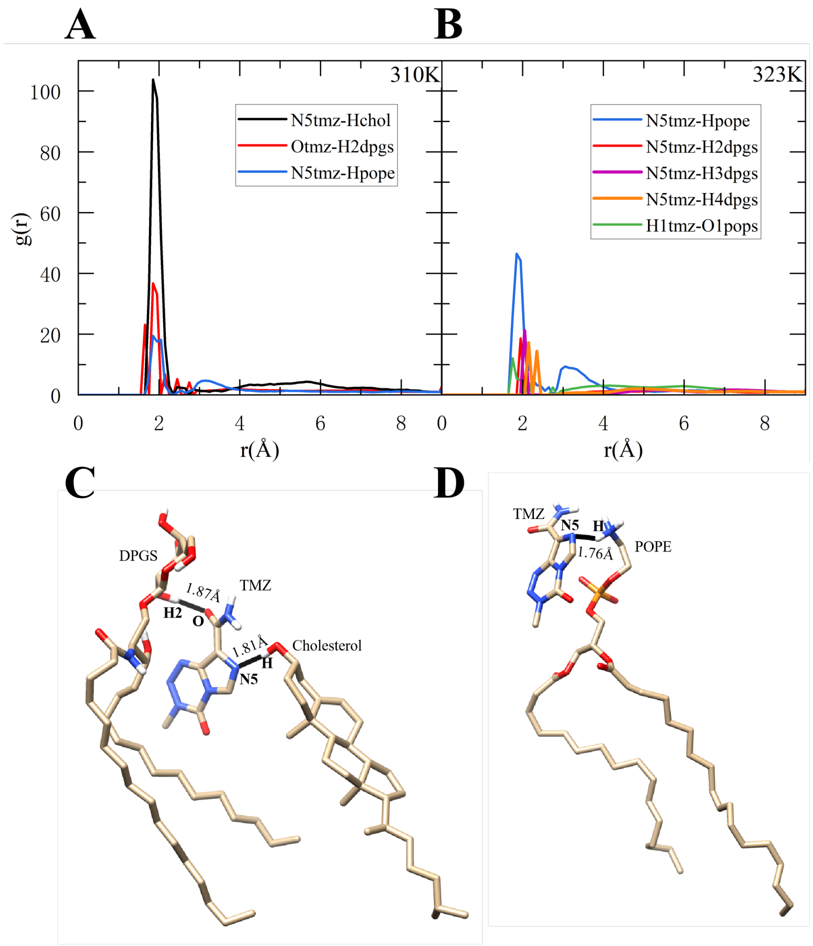


**Fig S6.** A,B: RDF for TMZ with various lipids while permeating the BPM. C,D refer to snapshots shows the interactions between TMZ and DPGS and CHOL at 310 K and that of TMZ and POPE at 323 K, respectively. Thick black lines indicate hydrogen bonds.
